# Validation and Application of a Protocol for the Extraction and Quantitative Analysis of Sphingomyelin in Erythrocyte Membranes of Patients with NAFLD

**DOI:** 10.1101/2021.04.16.440128

**Authors:** Charalampos Papadopoulos, Konstantinos Mimidis, Ioannis Tentes, Thaleia Tente, Konstantinos Anagnostopoulos

## Abstract

A set of constituents of the erythrocyte membrane lipidome has been proposed to serve as biomarkers for liver disease and acute coronary syndrome. In erythrocytes, sphingomyelin hydrolysis provides ceramide, a signaling lipid necessary for phosphatidylserine exposure and eryptosis. Phosphatidylserine exposure further amplifies hepatic inflammation and fibrosis during non-alcoholic fatty liver disease (NAFLD). In this study, we developed and applied a quantitative TLC for erythrocyte membrane sphingomyelin of NAFLD patients. We also compared 10 extraction methods for the isolation of sphingomyelin from erythrocytes. For quantitative TLC, lipids were separated in Silica gel 60 F254 using a mixture of chloroform/methanol/acetic acid/water (60/50/1/4) (v/v/v/v). The separated lipids were stained in a chamber containing iodine, and the intensity of each of the primary colors (red, green, blue) and the sum of the Red plus Green colors (R+G) was analyzed. The method was linear over a wide range of concentrations, presented acceptable precision (inter-day CV(%) 0.34, 0.006 and 0.44 for 2.5, 5.0 and 10 μg, respectively), good accuracy (recovery range 85.2-97.1%), and excellent limit of detection (0.137 μg/spot) and limit of quantification (0.41 μg/spot). Using this quantitation method, we compared various lipid extraction methods and found that lipid extraction with methanol led to higher yield of erythrocyte sphingomyelin (135.35±1.04% recovery, compared to the Folch method). Application of these methods showed that erythrocytes from NAFLD patients (9 men, 15 women, 57.95±11.11 years old) contained statistically significantly less sphingomyelin (829.82±511.60 vs 1892.08±606.25 μg/ml of packed erythrocytes) compared to healthy controls (4 men, 6 women, 39.3±15.55 years old).

## 1. Introduction

Lipid constituents of the erythrocyte membrane can be promising biomarkers for liver disease [1] and acute coronary syndrome [2]. The Phosphatidylcholine to Phosphatidylethanolamine ratio in the erythrocyte membrane has been correlated with the presence of Non-Alcoholic Fatty Liver Disease (NAFLD) [3] and resolution of hepatitis C [4]. Interestingly, during NAFLD, increased erythrocyte phosphatidylserine exposure leads to erythrocyte accumulation in the liver. Subsequently, erythrophagocytosis by Kupffer cells augments inflammation and oxidative stress [5]. Furthermore, we have recently found that red blood cells from NAFLD patients release increased Monocyte Chemoattractant Protein 1 (MCP1) chemokine and induce Tumor Necrosis Factor-α (TNF-α) release by macrophages in vitro (paper in preparation). In this context, membrane sphingomyelin hydrolysis could mediate a molecular communication between the inflammatory microenvironment in steatotic liver and phosphatidylserine exposure in erythrocytes. Dinkla et al [6] showed that during systemic inflammation, secreted sphingomyelinase induces sphingomyelin hydrolysis in red blood cells, further promoting Phosphatidylserine (PSer) exposure. This is of particular importance, since during NAFLD, serum acid sphingomyelinase displays a marked elevation [7], and acid sphingomyelinase is implicated in various molecular aspects of NAFLD pathology, such as apoptosis, endoplasmic reticulum stress, autophagy deregulation and fibrosis [8]. Thus, we hypothesized, that during NAFLD and in particular, during steatohepatitis, any changes in the sphingomyelin content in erythrocyte membrane can be indicative of a potential involvement of the erythrocyte in NAFLD pathology.

However, methods for lipid extraction can give variable results, optional to the choice of solvents. The most commonly used methods for lipid extraction are the Folch [9] and the Bligh and Dyer methods [10], which involve the treatment of erythrocytes with methanol/chloroform (in a ratio of 1:2) and a wash treatment with water, or the use of chloroform/methanol/water in ratio 1:2:0.8, respectively. Nevertheless, these methods have been found inappropriate for lipid extraction from erythrocytes [11]. This could be explained by the use of centrifugation, which results in an uneven distribution of lipids [12]. Furthermore, addition of water prior to lipid extraction has been found to decrease the extractability of certain lipids from erythrocytes [13].

Other researchers have recommended the use of different solvent systems for efficient lipid extraction from red blood cells. Rose and Oklander [11], after comparing the efficiency of different solvents for extracting cholesterol and total phospholipids, proposed the use of chloroform/isopropanol 7:11. Broekhuyse [14] proposed a system with multiple solvents. The inconvenience of multiple solvents and steps, led other researchers to develop methods requiring only a single step, involving either isopropanol or chloroform/isopropanol at a ratio of 1:1.5 [15–17]. Recently, Zhao and Xu [18], developed a rapid and efficient method for the analysis of phospholipids and lysophospholipids, which has not been evaluated for the analysis of erythrocyte sphingomyelin.

In this study we validated a quantitative thin layer chromatography method for sphingomyelin. Recently, Kerr et al [19] validated and applied a quantitative thin layer chromatography in combination with iodine staining and image analysis. In their work, they exploited the intensity of the bands, by analyzing the three primary colors (red, green, blue) in Microsoft Paint. After validation, we used this method for the validation of a quantitative thin layer chromatography for i) the comparison of ten lipid extraction methods for the selective isolation, and ii) quantitation of sphingomyelin in erythrocyte membranes of patients with NAFLD.

## 2. Experimental

### 2.1 Erythrocyte membrane isolation

Three milliliters of blood containing EDTA was centrifuged at 200 g for 10 minutes at 4°C. Plasma and buffy coat were removed. Then, 1 milliliter of erythrocyte pellet was washed with cold saline solution and centrifuged at 200 g for 10 minutes at 4°C. Erythrocytes were diluted 1:10 (v/v) with cold hemolysis solution (Tris 1 mM–NaCl 10 mM – EDTA 1 mM pH 7.2) and, after incubation for 30 minutes at 4°C with continuous shaking, they were centrifuged at 15000 rpm in a HERMLE 323 K centrifuge, 220.80 VO2 rotor, at 4°C for 16 minutes. The pellet was collected, and the last step was repeated for as many times as needed, until the final pellet was milky-white, signifying hemoglobin removal. Samples were kept at -80°C until analysis.

### 2.2 Lipid extraction

For all extraction methods, on 100 μL of erythrocyte hemoglobin-free membrane (ghost) the solvent system was added. Furthermore, for all solvent systems the sample/solvent volume ratio was adjusted at 1:12. Εvery extraction method was repeated twice and extraction efficiency is expressed as mean ± standard deviation of percent recovery by reference to an identical extraction by the Folch method (method number 1) which was set as 100% extraction efficiency. For the analysis of erythrocyte membrane sphingomyelin from patients, 50μl of ghost per sample was used and results were expressed as mg of sphingomyelin/ml of packed erythrocytes. The methods are described in Table 1.

**Table 1.**
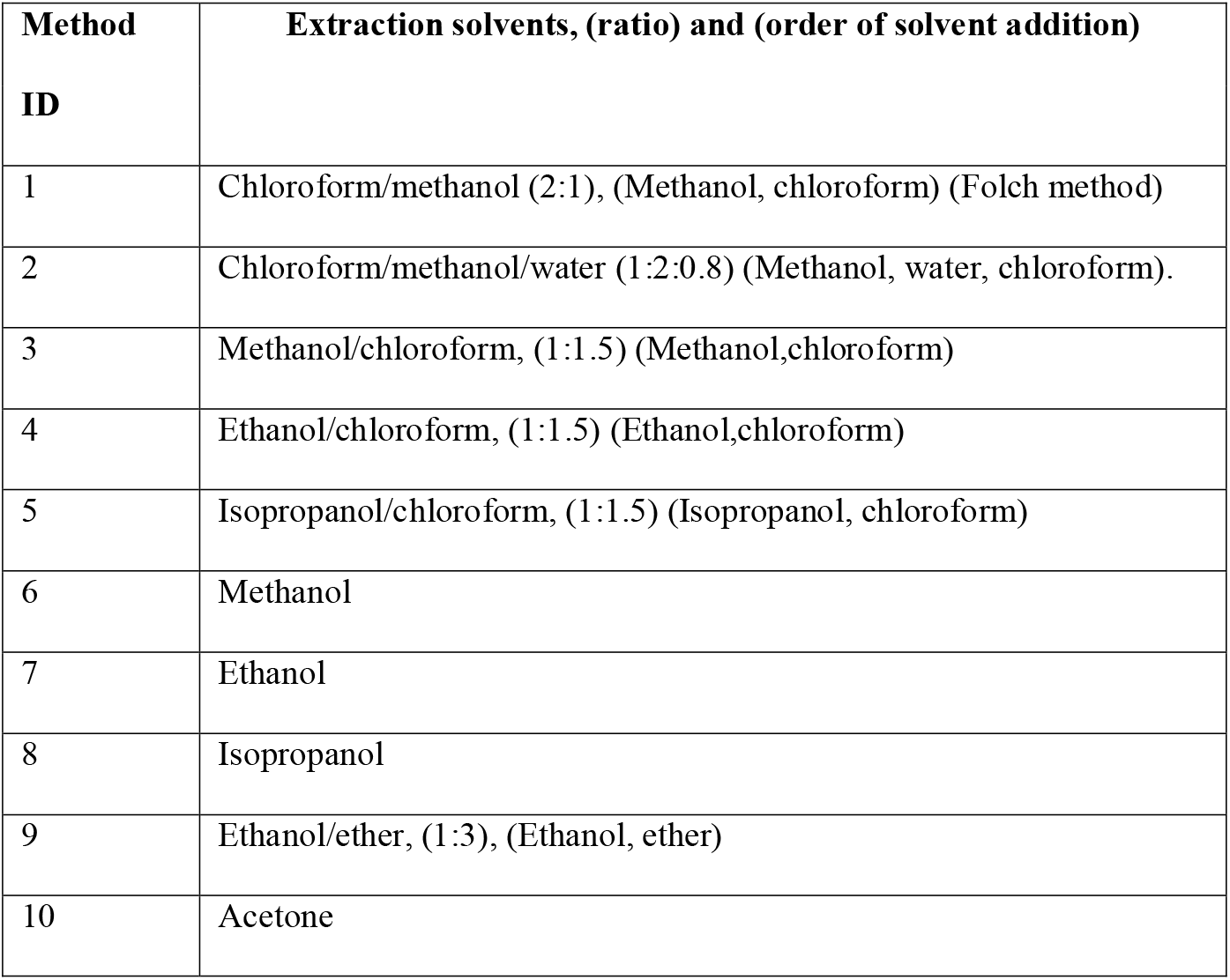
Methods of lipid extraction.

Samples were centrifugated at 5000g for 10 minutes, at 4°C. The organic phases were collected and evaporated at 70°C.

### 2.3 Lipid separation

Thin layer chromatographic analysis of erythrocyte membrane lipids was done on a 10×10 cm chromatographic plate (TLC Silica gel 60 F254 (Merck KGaA 64071 Darmstadt, Germany) using a mixture of chloroform/methanol/acetic acid/ water (60/50/1/4) (v/v/v/v). Before loading the samples, the plate was desiccated at 150° C for 10 minutes and pre-run with the developing solvent mixture as above. After sample separation, the plate was dried with hot air and then placed in a closed container of vaporized iodine (3.5 gr), placed facing the bottom of the container, for 30 minutes at room temperature. Lipids appeared as dark yellow bands against a lighter background. Lipids were identified by comparison with lipid standards, namely cholesterol, phosphatidyl-ethanolamine, phosphatidyl-inositol, phosphatidyl-serine, phosphatidyl-choline and sphingomyelin all purchased from Sigma-Aldrich (Munich, Germany).

### 2.4 Validation of a quantitative thin layer chromatography

The chromatograms were scanned using a scanner (Canon MX-475 with color scanning mode at a resolution of 300 dpi). The intensity of individual spots was estimated by using a new method for the quantitation of lipids, based on the study of Kerr et al [19]. The Microsoft Paint program was used, in which the images were opened. The color in the middle of the spots was picked using the “color picker” function of the program. Then, using the “edit colors” function, the RGB (red, green and blue) values of the component colors were determined and used for further analysis. In case there is no computer with the Windows operating system available, the same can be accomplished using the free open source image editor GIMP, which is available for Windows, OSX, Linux and other operating systems. In GIMP, the RGB values can be shown using the dockable pointer dialog (menu Windows – Dockable Dialogs – Pointer in GIMP 2.10).

For the determination of linearity, the analysis was repeated three times on three separate days at 5 concentrations (10, 5, 2.5, 1.25, 0.75, 0.5, 0.25 μg/spot). The color that gave the best results for linearity was selected for further experiments. The dynamic range was determined by analyzing 13 concentrations (200, 100, 50, 25, 10, 5, 2.5, 1.25, 0.75, 0.5, 0.25, 0.2 and 0.1 μg/spot). For the inter-day precision, the analysis was repeated three times on three consecutive days at three different concentrations (1, 0.5, 0.25 μg/spot). Precision was expressed as relative standard deviation (coefficient of variation, CV%). For the study of accuracy, we studied % recovery of 100%, 200% and 300% mass added in 2.5 μg/spot. The limit of quantification (LOQ) and limit of detection (LOD) were determined by repeating three times the analysis of three samples (0.25, 0.5, 0.75 μg/spot). Then the LOQ and LOD were expressed as 10^*^SD/slope and 3.3^*^SD/slope of the calibration curve, respectively [20].

### 2.5 Patients

24 patients (9 men, 15 women) and 10 healthy controls (5 men, 5 women) participated in our study. They were recruited by the 1st Pathology Clinic, Department of Medicine, Democritus University of Thrace, Alexandroupolis, Greece. All patients presented with hepatic steatosis according to ultrasonography. After exclusion of viral, alcoholic, drug and other causes, patients were evaluated by non-invasive biomarkers (NFS, FIB4, AST/ALT ratio). Their anthropometric characteristics are shown in the Results section 3.4. Our study was approved by the scientific council of the University Hospital of Alexandoupolis and the Ethics Committee after informed consent of the participants.

### 2.6 Statistical Analysis

Results are expressed as mean ± standard deviation, unless otherwise stated. Since the sample size was small, we used a Bayesian approach for statistical analysis, which is much more suited to provide meaningful results for small datasets. Bayesian analysis does not assume large sample sizes and smaller datasets can be analyzed while retaining statistical power and precision [21,22].

Statistical analysis and creation of figures (except Fig. 2) was done with the R programming language v. 3.6 [23]. Testing for difference between means was performed using the BEST package [24]. The results are reported as the probability P of the difference between the means of healthy subjects and NAFLD patients being less than zero, P (Healthy<NAFLD), or greater than zero, P (Healthy>NAFLD). This probability is more intuitive than the more convoluted meaning of the frequently used p value (the p value is the probability of observing the data, assuming the null hypothesis -no difference between means-is correct). Probabilities above 90% were considered statistically significant.

**Fig. 1.**
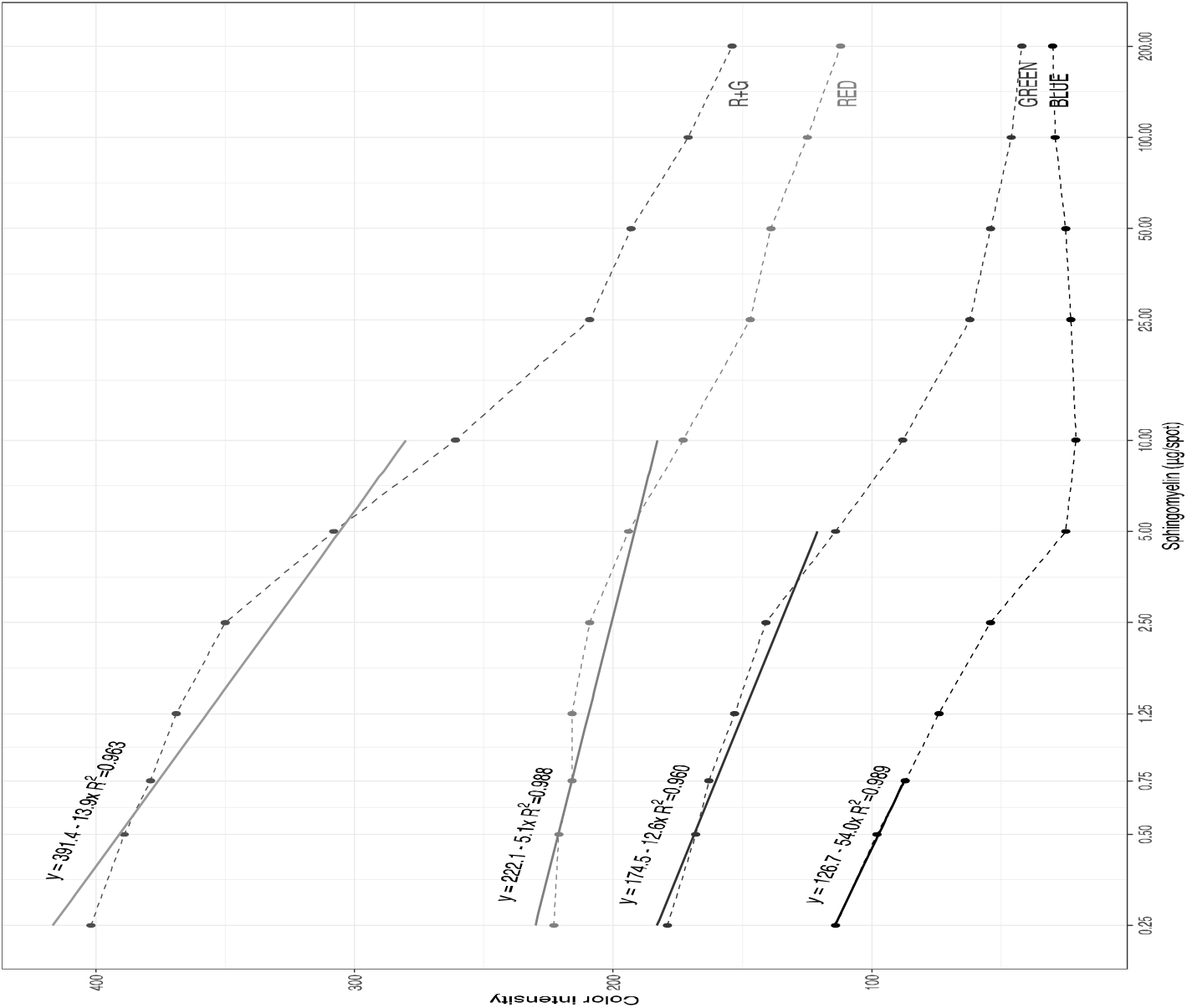
Response of color intensity versus sphingomyelin mass in TLC for the various color components (R+G=sum of Red plus Green color). The linear regression line for the linearity range is plotted (0.25-0.75 μg for Blue, 0.25-5.00 for Green, 0.25 – 10.00 for Red and R+G). The abscissa scale is log10

**Fig. 2.**
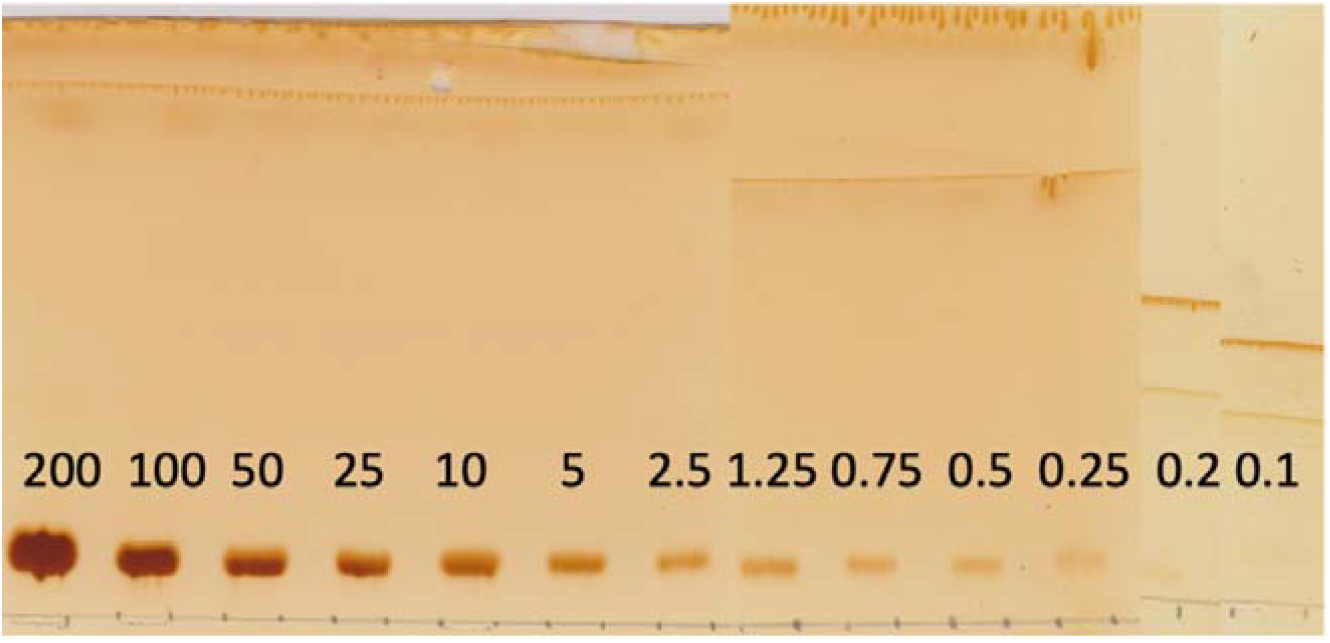
TLC image of sphingomyelin analysis for a range of masses (0.1-200 μg/spot)

## 3. Results and Discussion

### 3.1 Linearity of TLC

The method was linear over a wide range of concentrations (0.25-10 μg/spot), depending on the color component used, as shown in Figure 1. The R^2^ value was determined for each color for increasing ranges (0.25-0.50, 0.25-0.75, 0.25-1.25, 0.25-2.5, 0.25-5, 0.25-10 μg/spot). The best R^2^ value was used to determine the range of acceptable linearity. The best linearity was for the blue color, however the range of good linearity for it was limited (0.25-0.75 μg/spot). The green colour exhibited good linearity for the range 0.25-5.00 μg/spot. The red colour showed good linearity, as well as the sum of red plus green (R+G) colors, with a range 0f 0.25-10.00 μg/spot. The R+G was selected to be used for the following analyses, because the least squares lines of it showed a higher slope, which is indicative of higher sensitivity. The linear regression equations for the respective ranges are shown in Figure 1.

### 3.2 Dynamic range, specificity, sensitivity, precision and accuracy of the quantitative TLC

We next examined the range, limit of detection, limit of quantification, inter-day precision and accuracy of our method. The retention factors are shown in Table 2 and the performance characteristics in Table 3. The dynamic range of the methods was found to be 0.25-200 μg (Figure 2). However, in amounts greater than 10 μg/spot the linearity was lost, as described in the previous section. The limits of detection and quantification were 0.137 μg and 0.41 μg, respectively. The inter-day CV were 0.34, 0.006, 0.44 for 2.5, 5 and 10 μg/spot, respectively. The average accuracy, expressed as % recovery, was 91.0 with a range of 85.2 – 97.1.

**Table 2:** Retention factors of various membrane lipids in TLC

| LIPID | CHOL | PE | PI | PSer | PC | SM |
| --- | --- | --- | --- | --- | --- | --- |
| RF | 0.053 | 0.282 | 0.443 | 0.594 | 0.75 | 0.863 |

**Table 3:** Performance characteristics of the quantitative TLC for the analysis of sphingomyelin using the sum of the green plus red color.

| Performance characteristic | Value |
| --- | --- |
| Limit of detection ($\mu\text{g/spot}$) | 0.137 |
| Limit of quantification ($\mu\text{g/spot}$) | 0.41 |
| Range ($\mu\text{g/spot}$) | 0.2-200 |
| Inter-day CV (%) (2.5$\mu\text{g}$) | 0.34 |
| Inter-day CV (%) (5$\mu\text{g}$) | 0.06 |
| Inter-day CV (%) (10$\mu\text{g}$) | 0.44 |
| Recovery (%) (+100%) | 97.1±0.01 |
| Recovery (%) (+200%) | 90.8±0.007 |
| Recovery (%) (+300%) | 85.2±0.003 |

### 3.3 Extraction of sphingomyelin by 9 methods and comparison to the Folch method

For the purposes of our study, we sought to investigate which method extracts the highest amount of sphingomyelin from erythrocyte membranes. After comparing 9 methods (methods 2-10) with the Folch method used as reference (method number 1) which was calibrated as 100% extraction efficiency, we concluded that using methanol (method 6) as the extraction solvent method provides the best efficiency (135.35±1.04%). The results are shown in Figure 3.

**Fig. 3.**
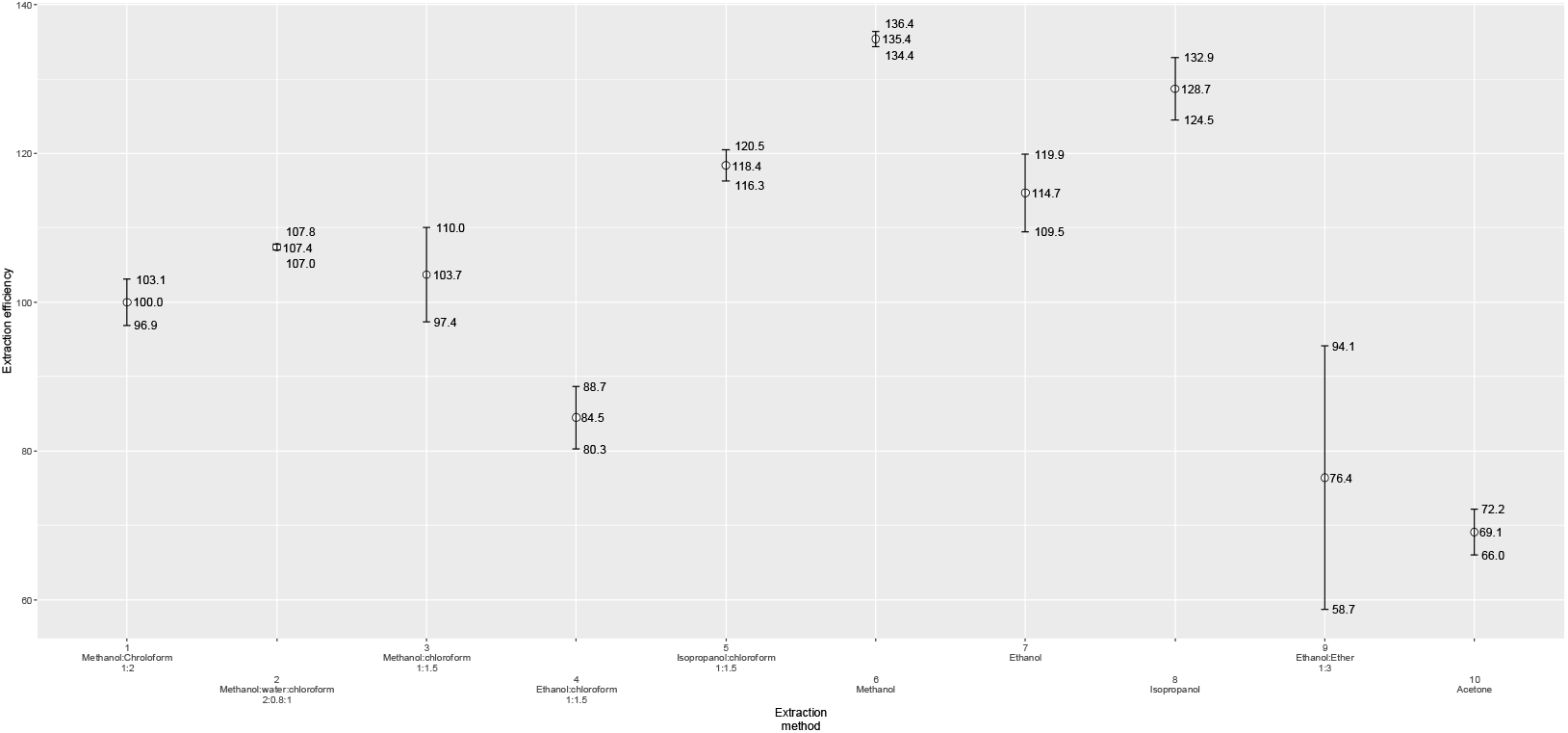
Extraction efficiency ± standard deviation for various lipid extraction methods. The solvents (where more than one) are mentioned by order of addition

Our study indicates that there are significant differences among the 10 different lipid extraction methods. The Folch method seems to be the best method for the extraction of cholesterol and total lipids. However, for the selective extraction of other lipids, other solvent systems should be used; for sphingomyelin, the method of Zhao and Xu (18), that also uses methanol in blood plasma, gives the best results, followed by the isopropanol method. For the purposes of our study, the methanol method (number 6) was selected for the analysis of erythrocyte membrane sphingomyelin in patients with NAFLD, because it exhibited the highest extraction efficiency.

### 3.4 Application of the quantitative TLC for the analysis of sphingomyelin of erythrocyte membranes from NAFLD patients and healthy controls

In order to test the developed method in a clinical setting, we measured sphingomyelin in erythrocyte membranes of NAFLD patients and healthy controls, as described in the Experimental section. The demographic and clinical characteristics of the study subjects are shown in Table 4, along with the test for statistical significance for differences between groups. It should be pointed out that the probability shown is not the p-value, but the probability that the groups are different in the Bayesian statistical framework, as described in the Statistical Analysis section 2.6.

**Table 4:** Characteristics of NAFLD patients and healthy controls.

| **PARAMETER** | **NAFLD Patients** | **Healthy controls** | **P(Healthy<NAFLD) or**<br>**P(Healthy>NAFLD)** |
| --- | --- | --- | --- |
| **AGE (YEARS)** | 58.0±11.1 | 39.3±15.5 | P(Healthy<NAFLD)=99.6% |
| **BMI** | 33.4±4.22 | 25.3±3.8 | P(Healthy<NAFLD)=100.0% |
| **AST (U/L)** | 46.5±19.3 | 21.7±8.5 | P(Healthy<NAFLD)=100.0% |
| **ALT (U/L)** | 80.0±66.8 | 21.9±6.9 | P(Healthy<NAFLD)=99.9% |
| **TRIGLYCERIDES (mg/dL)** | 224.1±159.9 | 152.3±18.1 | P(Healthy<NAFLD)=91.5% |
| **CHOLESTEROL (mg/dL)** | 196.9±46.9 | 154.2±14.2 | P(Healthy<NAFLD)=99.8% |
| **HDL-C (mg/dL)** | 45.4±7.3 | 65.1±14.0 | P(Healthy>NAFLD)=99.8% |
| **LDL-C (mg/dL)** | 109.8±42.6 | 58.6±7.50 | P(Healthy<NAFLD)=100.0% |
| **PLATELETS** | 223.3±63.4 | 240±53.5 | P(Healthy>NAFLD)=72.4% |
| **FIB4** | 1.90±1.28 | 0.77±0.35 | P(Healthy<NAFLD)=99.7% |
| **AST/ALT** | 0.91±0.50 | 0.97±0.16 | P(Healthy>NAFLD)=73.4% |

When sphingomyelin was measured using the procedure described above, erythrocyte membrane sphingomyelin of NALFD patients was significantly lower than healthy controls (829.82±511.60 vs 1892.08±606.25 μg/ml of packed erythrocytes, respectively) with P(NAFLD<Healthy)=100% (Figure 4).

**Fig. 4.**
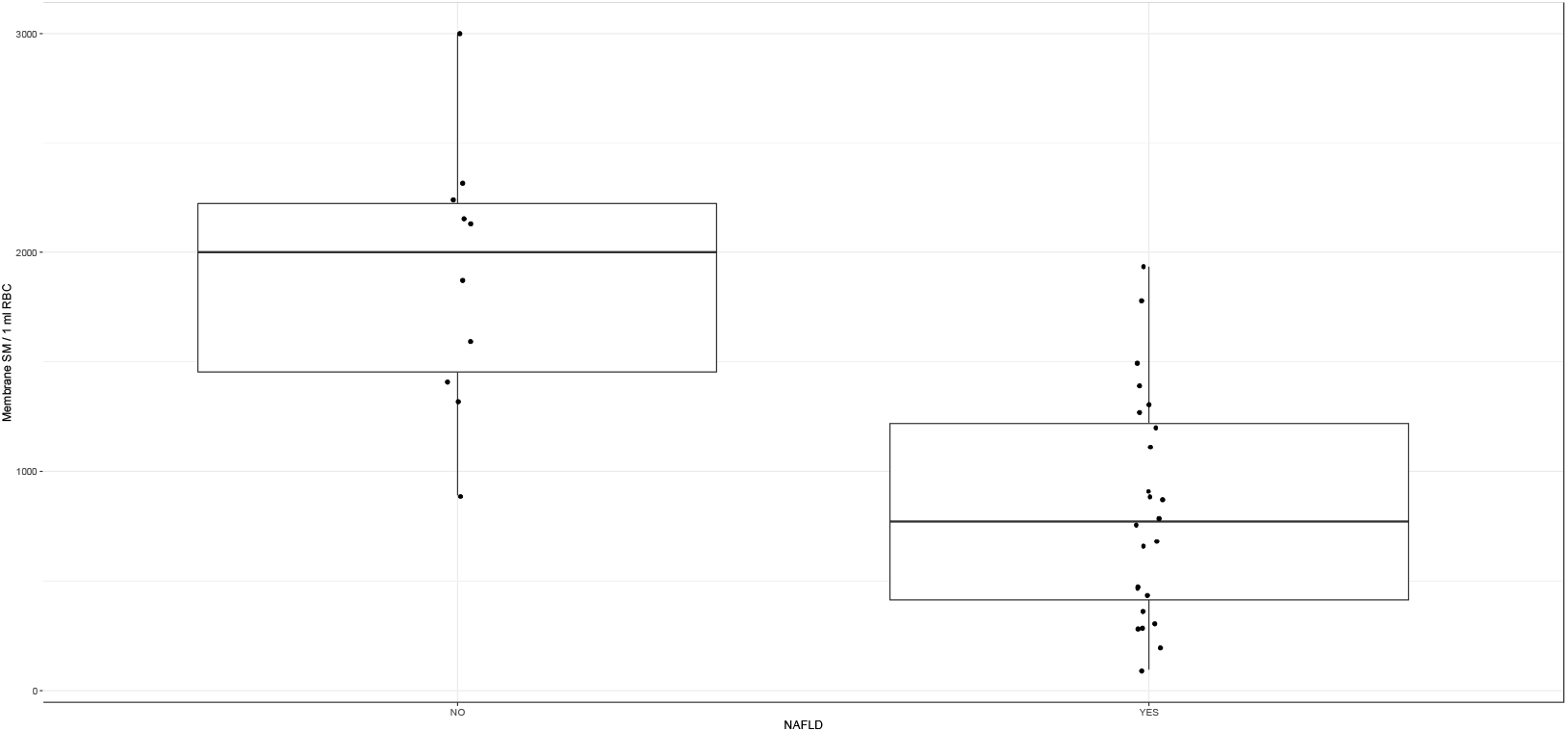
Boxplot of sphingomyelin levels in NAFLD patients and healthy controls

Our results are in agreement with the recent study of Chen et al [25]. In their study, the presence of both NAFLD and hyperlipidemia was correlated with low erythrocyte sphingomyelin. In our study, all patients presented hyperlipidemia (as evidenced by increased triglycerides, cholesterol and LDL and decreased HDL), and erythrocyte sphingomyelin was significantly lower than healthy controls. Since sphingomyelin hydrolysis provides ceramide, a signaling lipid necessary for phosphatidylserine exposure and eryptosis (erythrocyte death) [26] and phosphatidylserine exposure is further linked to the amplification of hepatic inflammation and fibrosis during NAFLD [5], our study merits further exploration.

## 4. Conclusion

In this study we describe a low-cost, precise, accurate and sensitive method for the analysis of sphingomyelin content in erythrocyte membranes. In addition, we have evaluated the recovery of sphingomyelin by 10 different extraction methods and found that methanol yields the highest amount. Finally, the application of our method for the quantitation of sphingomyelin of erythrocyte membranes of patients with NAFLD and healthy controls provided important results, with possible implications to the pathogenesis of the disease.

## Declarations

### Availability of data and material

Not applicable

### Ethics approval

The study was approved by the scientific council of the University Hospital of Alexandoupolis and the Ethics Committee after the informed consent of the participants.

### Consent to participate

Not applicable

### Consent for publication

Not applicable

### Code availability

Not applicable

### Authors contributions

Conceptualization: Charalampos Papadopoulos, Konstantinos Anagnostopoulos; Methodology: Charalampos Papadopoulos, Thaleia Tente; Formal analysis and investigation: Konstantinos Anagnostopoulos; Writing - original draft preparation: Charalampos Papadopoulos; Writing - review and editing: Konstantinos Anagnostopoulos; Funding acquisition: Ioannis Tentes; Resources: Konstantinos Mimidis; Supervision: Konstantinos Anagnostopoulos, Ioannis Tentes.

## References

1. Eapen CE, Ramakrishna B, Balasubramanian KA. Analyzing lipids in the liver & in the red blood cell membrane in liver diseases [Internet]. Vol. 147, Indian Journal of Medical Research. Indian Council of Medical Research; 2018. p. 334–6.

2. Tziakas DN, Kaski JC, Chalikias GK, Romero C, Fredericks S, Tentes IK, et al. Total Cholesterol Content of Erythrocyte Membranes Is Increased in Patients With Acute Coronary Syndrome. A New Marker of Clinical Instability? J Am Coll Cardiol. 2007;49(21):2081–9.

3. Arendt BM, Ma DWL, Simons B, Noureldin SA, Therapondos G, Guindi M, et al. Nonalcoholic fatty liver disease is associated with lower hepatic and erythrocyte ratios of phosphatidylcholine to phosphatidylethanolamine. Appl Physiol Nutr Metab. 2013;38(3):334–40.

4. Papadopoulos C, Panopoulou M, Mylopoulou T, Mimidis K, Tentes I, Anagnostopoulos K. Maedica-a Journal of Clinical Medicine Cholesterol and Phospholipid Distribution Pattern in the Erythrocyte Membrane of Patients with Hepatitis C and Severe Fibrosis, before and after Treatment with Direct Antiviral Agents: A pilot Study ErythrocytE Lipids in hEpatitis c. Maedica A J Clin Med. 2020;15(2):162–8.

5. Otogawa K, Kinoshita K, Fujii H, Sakabe M, Shiga R, Nakatani K, et al. Erythrophagocytosis by liver macrophages (Kupffer cells) promotes oxidative stress, inflammation, and fibrosis in a rabbit model of steatohepatitis: Implications for the pathogenesis of human nonalcoholic steatohepatitis. Am J Pathol. 2007;170(3):967–80.

6. Dinkla S, Wessels K, Verdurmen WPR, Tomelleri C, Cluitmans JCA, Fransen J, et al. Functional consequences of sphingomyelinase-induced changes in erythrocyte membrane structure. Cell Death Dis. 2012;3(10):e410–e410.

7. Grammatikos G, Mühle C, Ferreiros N, Schroeter S, Bogdanou D, Schwalm S, et al. Serum acid sphingomyelinase is upregulated in chronic hepatitis C infection and non alcoholic fatty liver disease. Biochim Biophys Acta - Mol Cell Biol Lipids. 2014;1841(7):1012–20.

8. Garcia-Ruiz C, Mato JM, Vance D, Kaplowitz N, Fernández-Checa JC. Acid sphingomyelinase-ceramide system in steatohepatitis: A novel target regulating multiple pathways [Internet]. Vol. 62, Journal of Hepatology. Elsevier; 2015. p. 219–33.

9. Folch J, Lees M, SLOANE STANLEY GH. A simple method for the isolation and purification of total lipides from animal tissues. J Biol Chem. 1957;226(1):497–509.

10. Reis A, Rudnitskaya A, Blackburn GJ, Fauzi NM, Pitt AR, Spickett CM. A comparison of five lipid extraction solvent systems for lipidomic studies of human LDL. J Lipid Res. 2013;54(7):1812–24.

11. Rose HG, Oklander M. IMPROVED PROCEDURE FOR THE EXTRACTION OF LIPIDS FROM HUMAN ERYTHROCYTES. J Lipid Res. 1965;6:428–31.

12. Wang W-Q, Gustafson’ A. Lipid determination from monophasic solvent mixtures: influence of uneven distribution of lipids after filtration and centrifugation [Internet].

13. Freyburger G, Heape A, Gin H, Boisseau M, Cassagne C. Decrease of lipid extractability of chloroform-methanol upon water addition to human erythrocytes. Anal Biochem. 1988;171(1):213–6.

14. Broekhuyse RM. Improved lipid extraction of erythrocytes. Clin Chim Acta. 1974;51(3):341–3.

15. Wang WQ, Gustafson A. Erythrocyte lipid extraction in alcohol-chloroform systems: a comparative study. Acta Chem Scand. 1994;48(9):753–8.

16. Peuchant E, Wolff R, Salles C, Jensen R. One-step extraction of human erythrocyte lipids allowing rapid determination of fatty acid composition. Anal Biochem. 1989;181(2):341–4.

17. Yan K-P, Hao J, Dan N, Chen C. A Simple and Improved Method for Extraction of Phospholipids from Hemoglobin Solutions. Artif Cells, Blood Substitutes, Biotechnol. 2008;36(1):19–33.

18. Zhao Z, Xu Y. An extremely simple method for extraction of lysophospholipids and phospholipids from blood samples. J Lipid Res. 2010;51(3):652–9.

19. Kerr E, West C, Kradtap Hartwell S. Quantitative TLC-Image Analysis of Urinary Creatinine Using Iodine Staining and RGB Values. J Chromatogr Sci. 2016;54(4):639–46.

20. Shrivastava A, Gupta V. Methods for the determination of limit of detection and limit of quantitation of the analytical methods. Chronicles Young Sci. 2011;2(1):21.

21. Lee SY, Song XY. Evaluation of the Bayesian and maximum likelihood approaches in analyzing structural equation models with small sample sizes. Vol. 39, Multivariate Behavioral Research. Lawrence Erlbaum Associates, Inc. ; 2004. p. 653–86.

22. Hox JJCM, Schoot R van de, Matthijsse S. How few countries will do? Comparative survey analysis from a Bayesian perspective. Surv Res Methods. 2012;6(2):87–93.

23. R: The R Project for Statistical Computing [Internet].

24. Kruschke JK. Bayesian estimation supersedes the t test. J Exp Psychol Gen. 2013;142(2):573–603.

25. Chen W, Shao S, Cai H, Han J, Guo T, Fu Y, et al. Comparison of Erythrocyte Membrane Lipid Profiles between NAFLD Patients with or without Hyperlipidemia. Int J Endocrinol. 2020;2020:1–12.

26. Lang E, Bissinger R, Gulbins E, Lang F. Ceramide in the regulation of eryptosis, the suicidal erythrocyte death. Vol. 20, Apoptosis. Springer New York LLC; 2015. p. 758–67.

